# Light management by algal aggregates in living photosynthetic hydrogels

**DOI:** 10.1101/2023.09.28.559714

**Authors:** Sing Teng Chua, Alyssa Smith, Swathi Murthy, Maria Murace, Han Yang, Michael Kühl, Pietro Cicuta, Alison G. Smith, Daniel Wangpraseurt, Silvia Vignolini

**Affiliations:** Yusuf Hamied Department of Chemistry, University of Cambridge, Cambridge CB2 1TN, United Kingdom; Marine Biology Section, Department of Biology, University of Copenhagen, Strandpromenaden 5, DK-3000 Helsingør, Denmark; School of Chemical Engineering, University of Chinese Academy of Sciences, 100040 Beijing, Beijing, China; Cavendish Laboratory, University of Cambridge, CB3 0HE Cambridge, United Kingdom; Department of Plant Sciences, University of Cambridge, Cambridge CB2 3EA, United Kingdom; Marine Biology Research Division, Scripps Institution of Oceanography, University of California San Diego, La Jolla, CA 92093-0205, USA; Department of Nanoengineering, University of California San Diego, La Jolla, CA 92093-0205, USA; Max Planck Institute of Colloids and Interfaces, 14476 Potsdam, Germany

**Keywords:** hydrogels, living materials, photosynthesis, optical modelling

## Abstract

Rapid progress in algal biotechnology has triggered a growing interest in hydrogel-encapsulated microalgal cultivation, especially for the engineering of functional photosynthetic materials and biomass production. An overlooked characteristic of gel-encapsulated cultures is the emergence of cell aggregates, which are the result of the mechanical confinement of the cells. Such aggregates have a dramatic effect on the light management of gel-encapsulated photobioreactors and hence strongly affect the photosynthetic outcome. In order to evaluate such an effect, we experimentally studied the optical response of hydrogels containing algal aggregates and developed optical simulations to study the resultant light intensity profiles. The simulations are validated experimentally via transmittance measurements using an integrating sphere and aggregate volume analysis with confocal microscopy. Specifically, the heterogeneous distribution of cell aggregates in a gel matrix can increase light penetration while alleviating photoinhibition compared to a flat biofilm. Finally, we demonstrate that light harvesting efficiency can be further enhanced with the introduction of scattering particles within the hydrogel matrix, leading to a four-fold increase in biomass growth. Our study, therefore, highlights a new strategy for the design of spatially efficient photosynthetic living materials that have important implications for the engineering of future algal cultivation systems.

**Significance Statement:** The ability to cultivate microalgae at scale efficiently would allow more sustainable production of food and food additives. However, efficient growth of microalgae requires optimised light conditions, which are usually challenging to obtain using biofilm cultivations mode: as the outer layer of cells are necessarily more exposed to incoming light than the inner layer, posing the problem of photoinhibition on the outer cells receiving too much light, and shading the ones below. Here we study both experimentally and numerically, how microalgae aggregates growing in the confinement of a hydrogel can provide an improved light distribution and therefore biomass growth is maximised. This study proposes new strategies on how to engineer future photobioreactors.

## Introduction

The escalating demand for novel materials with biomimetic functionalities has stimulated the development of so-called biohybrid systems, which are typically comprised of a soft hydrogel matrix hosting living cells that can perform various functions (1-4). Biohybrids incorporating photosynthetic organisms such as photosynthetic bacteria or microalgae have been proposed for diverse applications, ranging from chemical sensing (5-7), bioremediation (8), biotransformation (9), cell regeneration (10), bioelectronics (11, 12), hydrogen generation (13), and energy production by artificial leaves (14).

Algal-based biohybrid systems offer a highly effective platform for algal cultivation, mitigating several fundamental challenges associated with traditional photobioreactors where the algae are planktonic i.e. free in liquid suspension cultures (15-17). From notable improvements in space and water requirements (18, 19), to protection against contamination (20, 21) and environmental stress (22), studies have reported that biohybrids for algal cultivation show enhancement in photosynthesis (23) and growth rates (24), as well as increased production of secondary metabolites such as pigments and lipids (25, 26), compared to traditional methods. In addition, gel-encapsulated cultures provide distinct advantages for specific applications such as carbon capture (27), water treatment (28-30), and non-invasive metabolite harvesting (31), while preventing contamination of surrounding natural water systems and potential threats to native species (32).

In order for photosynthetic systems to function efficiently, it is crucial to achieve a homogeneous distribution of light throughout the entire material, while minimising both overexposure and self - shading (33, 34). Nonetheless, there is a lack of studies investigating light delivery in such hybrid living systems, or the underlying reasons for the observed improvements in performance. Most studies on light management in algae cultures have focused on liquid suspension cultures, where algal cells are homogeneously dispersed either freely in an aqueous phase (35-38) or within a gel matrix (39), rather than growing naturally into aggregates, which is instead what occurs when algae are encapsulated (23, 40). Thus, how aggregate formation impacts light propagation and photosynthetic efficiency in gel-immobilised algal cultures and photobioreactors remains a compelling and intriguing question that requires thorough investigation through optical modelling and experimental analysis.

In this study, we performed optical characterisation of the microalgal biomass confined within a gel matrix and studied how cell aggregation affects light management. More specifically, we measured the transmittance of light through agarose gel pads embedded with algal aggregates and compared the experimental values with predictions from Monte Carlo-based modelling of radiative transfer (41, 42). The simulated local scalar irradiance was coupled into a net photosynthetic rate (***P***_***net***_) model based on the Harrison model (43), providing insight into the expected ***P***_***net***_ in homogeneous algal biofilms and encapsulated algal aggregates in photobioreactor systems. We explored different incident irradiance levels and variable areal biomass densities, considering the impact of aggregate growth within the hydrogel. We also varied the cell seeding density and the scattering matrix on optical propagation efficiency, both computationally and experimentally. The results highlighted a fundamental difference in terms of light distribution between a biofilm and a gel-embedded distribution of algal aggregates, while also demonstrating a potential for further light-harvesting enhancement via modulating the scattering properties of the hosting matrix. Our findings have significant implications for the optimisation of light transport within immobilised algal cultures and the design of novel photobioreactors for microalgae cultivation and harvesting, which are now becoming increasingly important due to the growing commercial demand for sustainable food sources and additives (25, 44).

## Results

### Optical characterisation of immobilised biomass

The encapsulation of microalgae within a gel matrix inevitably leads to the formation of dense aggregates, primarily caused by cell division and the inability of the daughter cells to disperse within the physical confinement imposed by the encapsulating matrices (23, 27, 45, 46). This phenomenon is general, as the algae are physically constrained and it can be observed in a wide variety of biohybrids composed of different types of algae and hydrogels (**Figure 1A, 1B, and 1C**).

**Figure 1.**
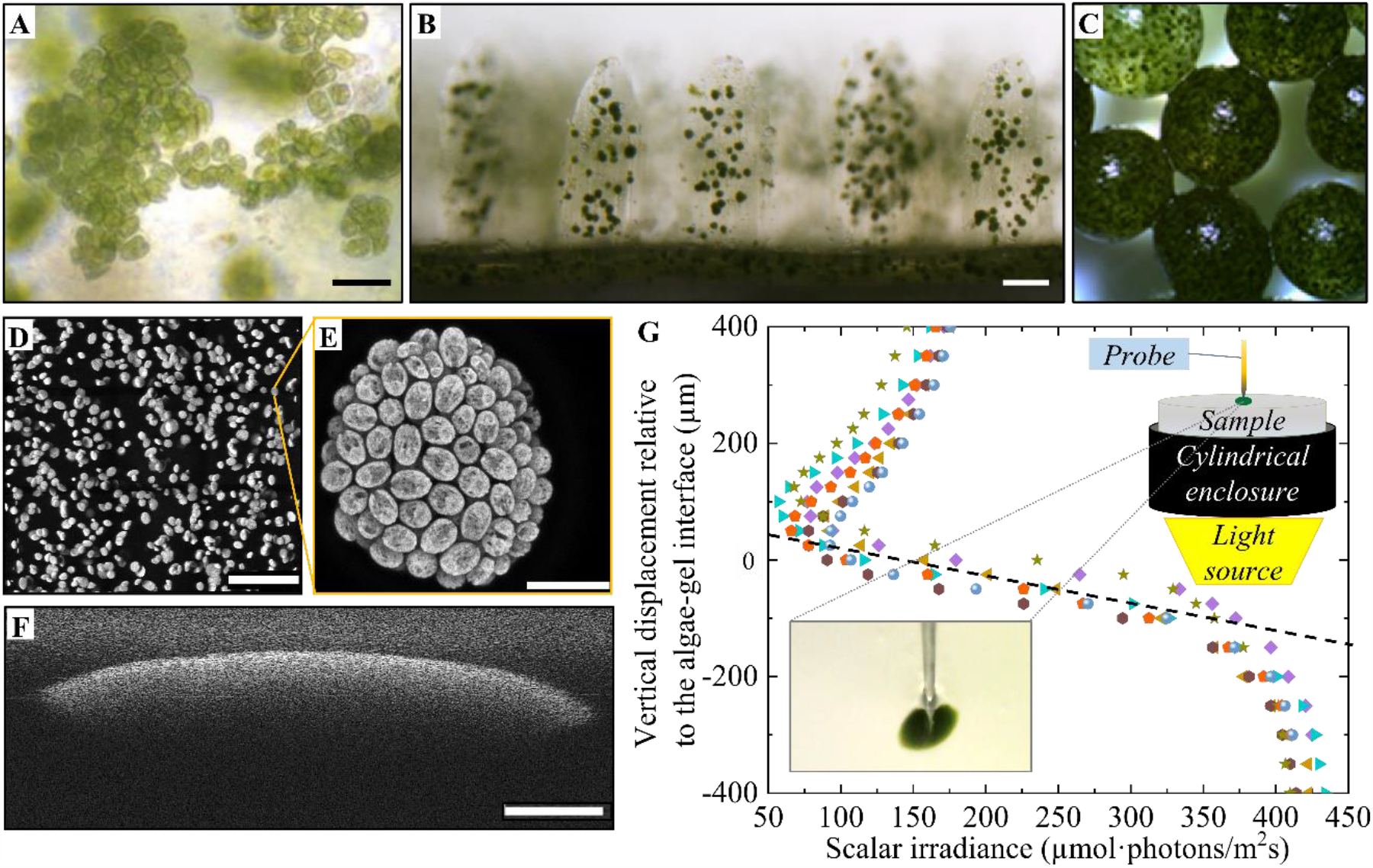
Cell aggregation resulting from the gel-immobilised culture of microalgae: (A) Isolated colonies of Platymonas sp. in silk-based hydrogel (scale bar = 10 μm) adapted with permission from (46) under Copyright 2023 American Chemical Society; (B) Aggregates of Marinichlorella kaistiae KAS603 in a 3D-printed bionic coral (scale bar = 100 μm) reprinted from (23) under Creative Commons CC-BY; (C) Tight clusters of Chlorella vulgaris encapsulated within sodium alginate (reprinted from (27) under Creative Commons CC-BY); (D) Confocal imaging (mean Z-stack projection) of chlorophyll autofluorescence emitted from Chlamydomonas reinhardtii aggregates after 7 days of growth within an agarose hydrogel (scale bar = 1 mm); (E) Close-up maximum intensity projection of Z-stack through a single aggregate (scale bar = 25 μm); (F) Cross-sectional view of an immobilised algal aggregate imaged with an optical coherence tomography system (scale bar = 500 μm); (G) Microsensor profile of photon scalar irradiance (400-700 nm) measured across individual algal aggregates, where the origin was set to be the upper interface between the aggregate and gel matrix. The top-right inset shows the schematic illustration of the microsensor measurement set-up. The bottom-left inset illustrates a close-up photograph of the microsensor tip penetrating an isolated algal aggregate.

In this study, to evaluate the capability of light management within such biohybrids, we considered model systems composed of microalgae *Chlamydomonas reinhardtii* encapsulated in agarose gel. However, many of the considerations we presented can be extended to other types of biohybrids. The first step for modelling realistic experimental conditions was to evaluate the precise shape of the aggregates and their scattering capability. To achieve this, we exploited both confocal microscopy and optical coherence tomography (OCT) techniques, revealing that the aggregates developed a lenticular, almost spherical shape (**Figure 1D, 1E, and 1F**). To conduct optical simulations, we used a simplified model where the gel-encapsulated culture was represented as a random arrangement of spherical algal aggregates embedded within the gel matrix. This approximation was made based on the observation that the oblateness of the aggregates had minimal effects on the optical attenuation results (**SI Figure 1**).

To evaluate the scattering parameters, we performed OCT with 930 nm light on isolated microalgal aggregates to estimate the scattering coefficient μ_*s*_ and anisotropy factor *g* within the confined biomass (**Figure 1F**; **SI Figure 2**). The empirical fitting of the backscattered intensity suggested that the values of μ_*s*_and *g* for 930 nm within individual algal aggregates were 1000 ± 100 cm^-1^ and 0.99, respectively. More details on how these values were extrapolated from the raw data is reported in the SI.

To assess the extent of light attenuation, the variation of spectral scalar irradiance was measured within individual aggregates, as a function of depth beneath the algae-gel interface. Considering the integrated spectral range from 400 to 700nm, which corresponds to the photosynthetically active radiation (PAR), the value of the scalar extinction coefficient μ_*ext,PAR*_ was calculated from the slope of light attenuation (**Figure 1G**) and found to be 180 ± 20 cm^-1^ based on the following equation (47).

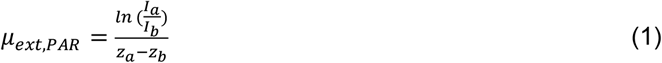

It is known that the scalar irradiance attenuation is equal to the absorption coefficient divided by average cosine of all the incident photons at medium interface (48). Considering that the algal biomass was highly forward scattering, assuming minimal wavelength dispersion of scattering anisotropy, the attenuation coefficient measured from scalar irradiance was close to the absorption coefficient, within the percentage uncertainty of measurement. Hence, we determined the absorption coefficient μ_*a*_ to be 180 ± 20 cm^-1^. Notably, this estimation aligns closely with the reported values from microalgal biofilms (49), suggesting that the extent of optical attenuation within isolated individual aggregates is comparable to that observed in dense biofilms.

### Simulation of light propagation through gel-immobilised aggregates

In order to simulate the effect of light propagation through gel-immobilised aggregates, it is important to consider that the system was not static, but was evolving continuously, with gel-encapsulated colonies developing into aggregates from individual algal cells that underwent cell growth and division. Therefore, different growth stages were simulated with different aggregate sizes under constant aggregate positions and numbers (**Figures 2A and 2B**). As expected, we observed a reduction in normalised scalar irradiance with the simulation depth, indicating greater light attenuation through absorption and scattering. The growth of the aggregate gave rise to greater attenuation of light with depth, due to the increased number of algae absorbing light (**Figure 2B**). It is also important to notice that the intensity plateaued on a non-zero value, indicating that light was also scattered more efficiently and not fully absorbed.

**Figure 2.**
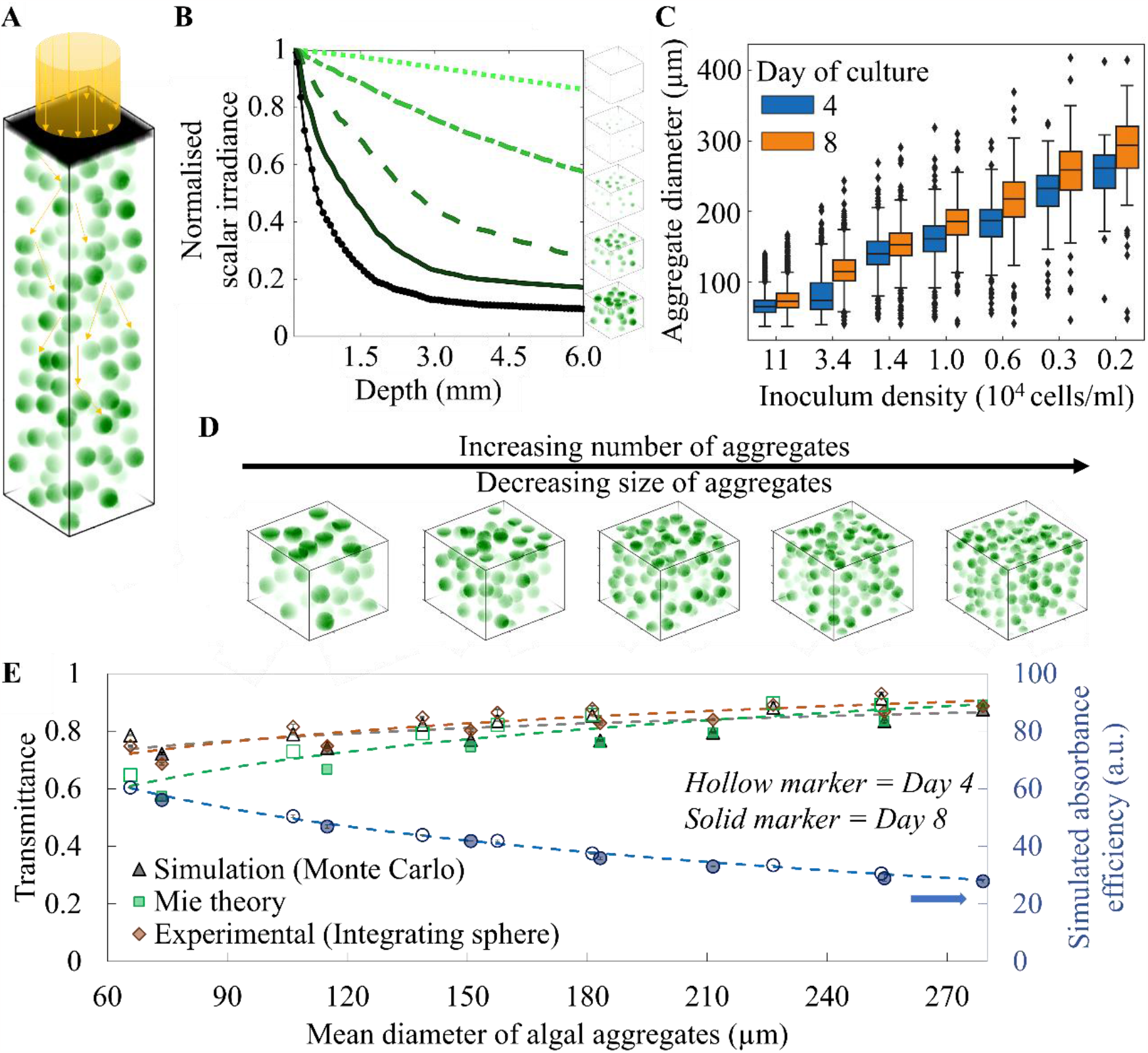
(A) Schematic of modelled configuration in the Monte Carlo simulation showing the downwelling beam illuminating upon microalgal aggregates represented as spheres randomly distributed within the agarose hydrogel, ranging from 10 to 100 μm in diameter; (B) Simulated attenuation of scalar irradiance normalised against the incident value among algae aggregates of different sizes but constant density; (C) Quantification of algal aggregate size distribution with varied inoculum density using confocal microscopy on day 4 and day 8 during cultivation; (D) Illustrative example of constant algal density with fewer large algal aggregates and a greater number of small algal aggregates; (E) Comparison of total transmittance measured from the gel samples of varied algal aggregate size distributions on day 4 (hollow marker) and on day8 (solid marker) using the integrating sphere against the Monte Carlo simulations and the analytical calculations with Mie’s theory and Beer-Lambert’s law. The secondary axis illustrates the simulated values of the overall absorbance efficiency, marked by blue circles.

To validate the simulation results of light transmission experimentally, we studied the effect of aggregate size and density of *C. reinhardtii* using integrating sphere measurement and quantitative analysis of aggregates through confocal imaging (**Figure 2C**). To prepare samples with algal aggregates of varying size and density distribution, we inoculated the culture with varying cell densities. Specifically, we observed that a lower inoculum density led to the formation of sparser but larger algal aggregates, while a larger inoculum density led to smaller aggregates (**Figures 2C and 2D**). **Figure 2C** also shows that independently from the inoculation density, the size of the aggregates increased from day 4 to day 8, indicating that the cells were growing and dividing actively during the culture period.

Transmittance spectra of gel-encapsulated cultures for all the different inoculation conditions were measured after 4 and 8 days of growth (**Figure 2E**). Here we observed that light attenuation was more pronounced when aggregates were smaller and denser, while larger but more aggregates gave a lower light attenuation. To evaluate the effective light absorbance for these different conditions, we performed Monte Carlo simulations. To develop the simulation model, we considered numerous algal aggregates encapsulated within gel culture, according to the spatial distribution and size variance of the algal aggregates obtained with confocal microscopy. As a result, we observed that the formation of sparser but larger algal aggregates contributes to higher optical transmittance, as supported by the numerical simulation and Mie theory prediction. The numerical simulation and experimental results deviated slightly from the theoretical prediction, which computed the effective attenuation coefficient using Mie’s theory and calculated the overall light transmission with Beer-Lambert’s law, as the assumption of homogeneous media implied in Beer-Lambert’s law did not hold and could not account for localised shading of light within individual aggregates. It provided less effective light absorbance per unit biomass volume, as expressed in our simulations by the lower percentage of the incident photon energy deposited per unit voxel (**Figure 2E**).

### Comparison of light transmission and utilisation within the algal biofilm and gel-immobilised algae

In contrast to planktonic cultures, the biofilm and gel-immobilised cultures share similarities in terms of higher biomass production and harvesting simplicity (1, 50). We, therefore, compared the capacity of the total areal biomass production per volume in these two configurations.

We modelled the gel-immobilised cultures and biofilm with an equal areal biomass density so that the total amount of algae and surface area of illumination were kept constant in all scenarios (**Figure 3A**). The simulation model considered lateral light loss with photons escaping in all directions, i.e., top, bottom, and four sides of the simulation volume. The lateral dimension was chosen to be at least twice that of the simulation depth, to minimise simulation artifacts in the form of edge effects. For a better comparison across varied areal biomass densities, the illuminated area of the simulation volume was kept constant at z = 1 mm and x = y = 2 mm.

**Figure 3.**
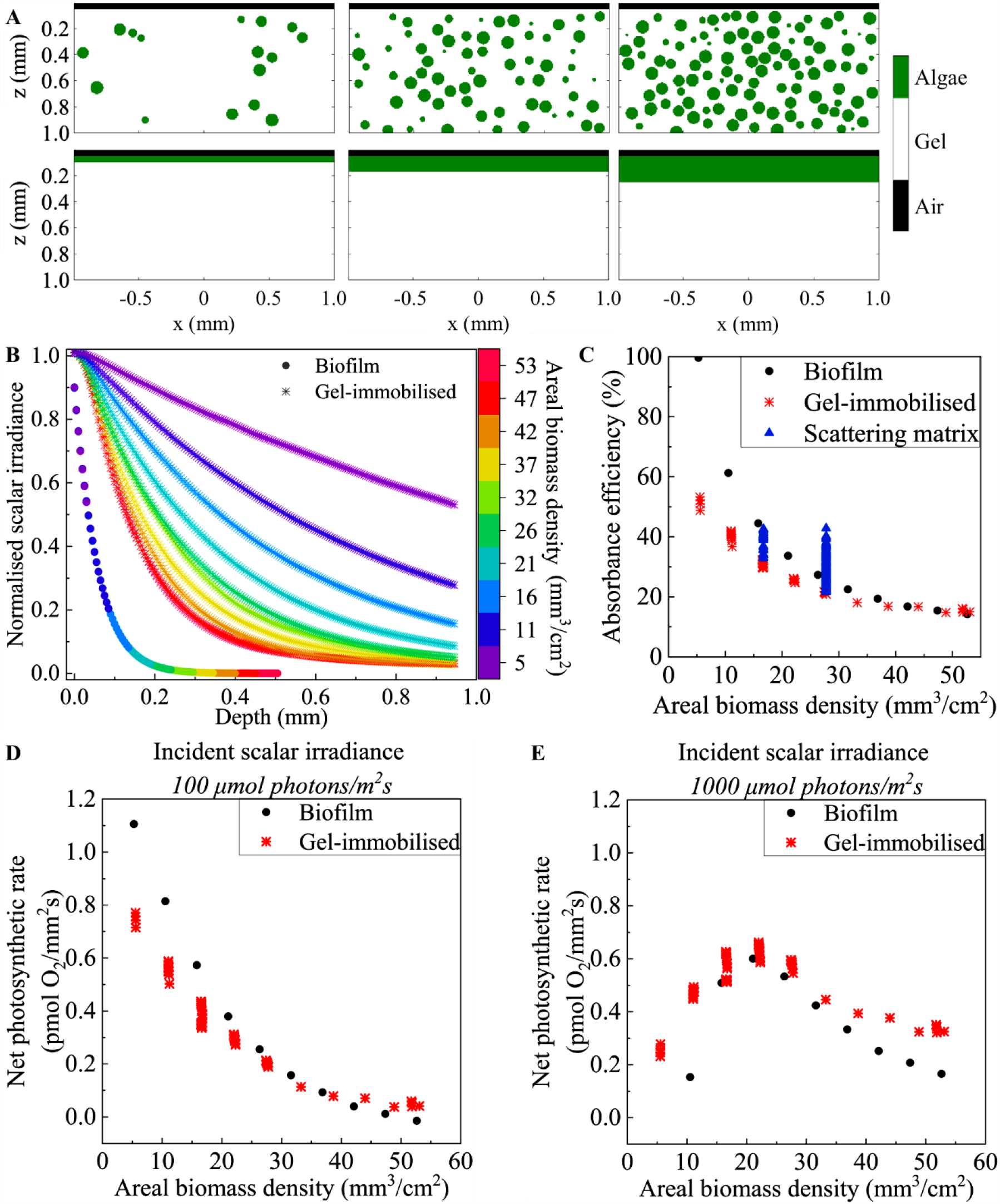
(A) Schematic cross sections of gel-immobilised culture (top) and biofilm (bottom) of equal areal biomass density in simulation; (B) 3D Monte Carlo simulation of scalar irradiance attenuation across biofilm (represented by ●) of varied thicknesses and across the gel culture (represented by *) with a mean aggregate radius of 50 μm but different aggregate densities, corresponding to varying areal biomass densities as represented by the colour scale; (C) calculation of the total light absorbance efficiency and the net photosynthetic rate (**P**_**net**_) by coupling the experimental light response curve to the simulated variation of scalar irradiance among algal aggregates and through biofilm of different areal biomass densities under an incident scalar irradiance of (D) 100 and (E) 1000 μmol photons m^-2^ s^-1^.

However, different biofilm thicknesses were also explored from z = 0.5 mm to 5 mm, keeping the constant area of illumination (x = y = 2 mm) and same areal density of algal biomass in gels compared to biofilms. A higher areal biomass density corresponded to a thicker biofilm and a denser aggregate distribution in gel immobilisation, and *vice versa*.

Owing to the dense distribution of absorbers (algae) in a biofilm, the normalised scalar irradiance approached zero at a depth of 200 μm (**Figure 3B**), consistent with previous reports, where biofilm growth was typically limited to a thickness of 200 μm to 300 μm (49, 51, 52). In contrast, the gel-encapsulated system exhibited much less light attenuation, resulting in similar transmittance output (**Figure 3B**). Given high enough areal biomass density, the gel-immobilised algal aggregates eventually exhibited light attenuation similar to that in biofilm with self-shading among aggregates.

At very high biomass densities, the aggregates tended to overlap and coalesce, forming a network of interconnected biomass, that could be observed both in simulations and practical experiments (**SI Figure 3**).

The understanding of light propagation and absorbance alone was, however, insufficient to characterise biomass production as the photosynthetic rate is not linear with the amount of light absorbed. To correlate the effective scalar irradiance to microalgae photosynthetic activity across the simulation volume depth, we coupled the optical simulation outcome to an experimentally determined light response curve fitted to an empirical model (**SI Figure 4**), taking the effect of photoinhibition into account. Specifically, we used the Harrison model (43) in estimating the net photosynthetic rate ***P***_***net***_, based on a known value of local scalar irradiance ***I***:

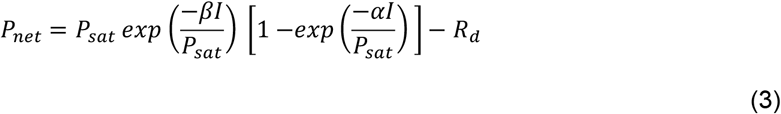

where *P*_*sat*_ indicates the maximum ***P***_***net***_ while *α* and *β* represent the light-limited initial slope and photoinhibition constant respectively. The rate of dark respiration is represented by ***R***_***d***_.

***P***_***net***_ was computed for all configurations using a range of incident irradiance levels from 100 to 2000 (natural sunlight) μmol photons m^-2^ s^-1^ (**SI Figure 5**). Comparisons were made between the biofilm and aggregate system by integrating ***P***_***net***_ across all depths, before normalising it by biomass density.

At low incident irradiance (100 μmol photon m^-2^ s^-1^), the integrated ***P***_***net***_ followed the trend of light absorbance efficiency in general (**Figure 3C**), as it fell in the light-limiting regime, with higher biomass density resulting in lower ***P***_***net***_ due to greater light attenuation (**Figure 3D**). The biofilm generated higher ***P***_***net***_ compared to the gel-immobilised system up to the thickness threshold where the light was attenuated completely in the biofilm. Beyond this threshold, around 35 mm^3^/cm^2^ in areal biomass density, the gel-immobilised configuration exhibited slightly higher ***P***_***net***_, as the bottom of the biofilm was heavily shaded, while the gel-immobilised system still had some light reaching the aggregates at the bottom, allowing for moderate photosynthetic activity.

Under high incident irradiance (1000 μmol photon m^-2^ s^-1^), photoinhibition became significant, as predicted by the Harrison model (**SI Figure 6**). Given high absorbance efficiency and less shielding effect at low areal biomass density, more cells were photo-inhibited, giving rise to lower ***P***_***net***_. On the other hand, with the increase of areal biomass density, the extent of self-shading increased, lowering the degree of photoinhibition, until the point of light saturation for maximal photosynthetic capacity. With further self-shading, part of the shielded culture became light-limited (**Figure 3E**). While the top layer of the biomass may be photo-inhibited, the shaded biomass would be exposed to the optimal light level. Overall, the gel-immobilised system exhibited higher ***P***_***net***_, indicating a lower degree of photoinhibition, as compared to the biofilm system.

### Performance optimisation with scattering matrix

To increase the absorbance efficiency, and therefore increase the probability of interaction between photons and algal aggregates, we explored the effect of incorporating scattering elements within the matrix. While a heterogeneous biomass distribution would enable the transmission of some photons among cell aggregates without interaction with the biomass, a scattering matrix with a wide angular range of scattering directions could redirect these unabsorbed photons to interact with the algal aggregates, effectively improving the probability of light absorbance. Experimentally, such enhancement of scattering efficiency of a matrix can be achieved by incorporating scattering particles into the gel matrix. **Figure 3C** shows that increasing the scattering efficiency in the surrounding matrix increased the effective absorbance per aggregate beyond that of the biofilm at high areal biomass density. A key parameter to consider when adding scattering element in the matrix is their filling fraction and distribution. The simplest approach is to use a uniform dispersion of scatterers throughout the gel matrix, with overall concentration per unit volume of scatterers, which scaled with the scattering coefficient in the unit of reciprocal length. Alternatively, further optimisation could be achieved by concentrating scatterers at different depths of the gel matrix, as illustrated in **Figure 4A**.

**Figure 4.**
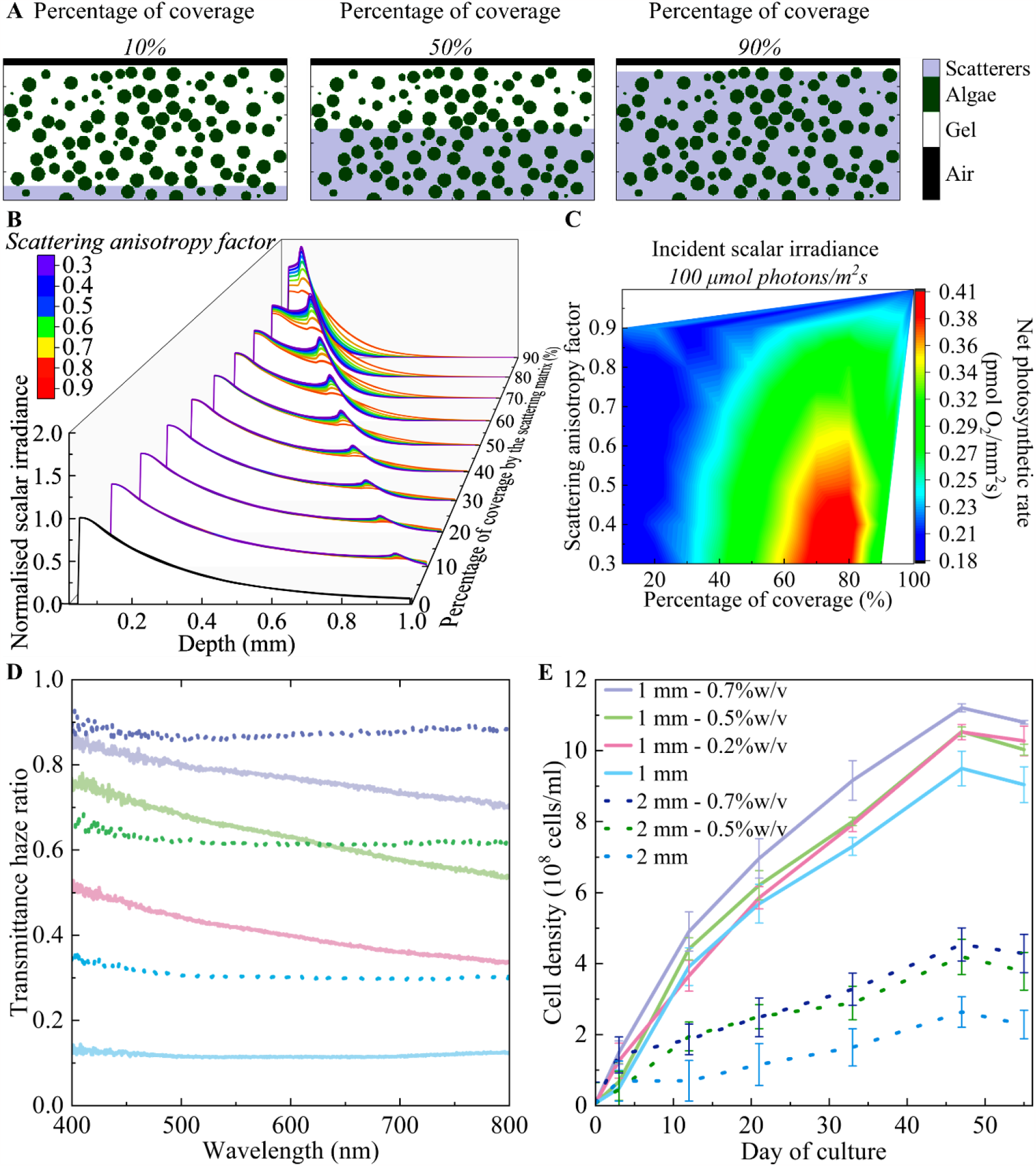
(A) Schematic cross-sections of varied scattering matrix coverage in percentage, with the scatterers being concentrated near the bottom boundary; (B) 3D Monte Carlo simulation of light attenuation across the gel culture with a mean aggregate radius of 50 μm and an areal biomass density of 27 mm^3^/cm^3^, but varying scattering matrix coverage and scattering anisotropy represented by the colour scale; Calculation of the net photosynthetic rate (**P**_**net**_) by coupling the experimental light response curve to the simulated variation of normalised scalar irradiance among algal aggregates within different scattering matrix configurations under an incident scalar irradiance of (C) 100 μmol photons m^-2^ s^-1^; (D) The effect of CMP doping on the transmittance haze across 1 mm-(solid line) and 2 mm-(dotted line) thick agarose hydrogels, across different weight percentage of embedded CMP; (E) The cell growth curve in 1 mm-(solid line) and 2 mm-(dotted line) thick hydrogels, colour matched to the CMP doping level it represents.

Our simulations showed how adding scattering particles to the gel matrix with different anisotropy factors and the percentage of coverage affected the light distribution within the matrix (**Figure 4B**). In contrast to a smooth exponential decay of normalised scalar irradiance in a material without a scattering matrix (represented by the black line), the interface between a normal gel matrix and a scattering matrix created a ripple of intensity profile, as a result of enhanced backscattering. The degree of this enhancement was influenced by the anisotropy factor of the scattering particles, with a lower anisotropy factor resulting in more pronounced backscattering effects. We note that following the peak of enhanced scalar irradiance at the medium-gel interface, light availability in deeper parts of the gel was diminished, as the scattered photons were either absorbed by the algae or escaped from the upper interface. The associated shifts in the profile of normalised scalar irradiance affected the overall absorbance efficiency and hence the integrated ***P***_***net***_ of the entire biomass volume, depending on the incident irradiance (**Figure 4C, SI Figure 7**).

In the gel-immobilised system, mutual shading among the cells in the aggregates resulted in the concentration of photons in the upper region, as light is attenuated with depth. Hence, under low incidence irradiance, increasing the availability and absorbance of photons in the upper region significantly enhanced the overall photosynthetic efficiency, particularly with a higher percentage of scatterer coverage and lower anisotropy factor (**Figure 4C**).

In addition, the distribution of the scatterers is important. Excessive backscattering especially in the upper region of the gel caused photoinhibition and reduced photon transfer to the lower regions when the scatterer coverage exceeded 70%. This was demonstrated in the case of 90% scatterer coverage, where the maximum scalar irradiance reached up to twice the incident irradiance. (**Figure 4B**). Moreover, since under high incident irradiance, the algae in the upper region of the gel were subjected to photoinhibition without any scattering presence, scattering enhancement in the upper area of the gel would lower the overall net photosynthetic efficiency. In contrast, the scattering enhancement was optimal in the light-limiting regime, namely the middle zone (40% to 50% in depth) after some degree of light attenuation from mutual shading (**SI Figure 7a**). With an even higher irradiance (1000 μmol photon m^-2^ s^-1^), the optimal zone for scattering matrix coverage reduced to the lower region of 20% to 40% in depth, as the light-limiting regime was shifted further downwards (**SI Figure 7b**). Under high irradiance, if the scattering interface was in the upper region with 80% coverage in depth, the ***P***_***net***_ decreased with a lower anisotropy factor as photoinhibition was aggravated with higher photon interaction in the light-saturated region (**SI Figure 7c and 7d**).

As a proof-of-concept experiment, agarose hydrogels were embedded uniformly with cellulose microparticles (CMP) (53) to increase light scattering within the matrix. CMPs were chosen as scattering particles for their optimized scattering abilities and their biocompatible characteristic. As evident in **SI Figure 8** the addition of CMP resulted in increased opacity of the hydrogel, owing to higher haze ratio of the haze transmittance to the total transmittance with higher proportions of CMP (**Figure 4D; SI Figure 9**). CMP doping introduced significant light scattering within the hydrogel matrix. Within the 1 mm-thick hydrogels, 0.7%w/v CMP doping caused 75%-85% of the transmitted light to be scattered, compared to approximately 15% in standard hydrogels without CMP (0%w/v). When comparing the spectra of the 2 mm hydrogels, 0.7%w/v CMP doping scattered approximately 90% of the transmitted light. This demonstrates that the addition of CMP altered the light profile throughout the hydrogels, thereby influencing the growth of immobilised microalgae (**Figure 4E**).

Within the 1 mm-thick agarose pad, in comparison to those without CMP, those containing 0.7%w/v CMP showed a higher biomass throughout the entire growth curve, showing an approximate 50% increase in cell numbers after 10 days. Additionally, there was cumulatively improved growth in the agarose with 0.7%w/v CMP, rather than an initial increase, which was then maintained. In 2 mm-thick agarose pads, both 0.5%w/v and 0.7%w/v CMP doping resulted in a higher degree of growth enhancement, with approximately 100% increase in cell numbers after 10 days (**Figure 4E**). Comparing the cell density alone, the 1 mm-thick hydrogels were much more productive than the 2 mm-thick hydrogels, with or without scattering, as a result of limited gaseous and mass exchange with increasing hydrogel thickness.

## Discussion

By comparing the light management capabilities of biofilms and gel-immobilised cultures, we have been able to conclude that gel-immobilised algal cultures have the potential to reach a higher areal biomass density compared to flat, homogeneous algal biofilms. Our results suggest that the formation of cell aggregates upon gel immobilisation is crucial as it reduces the probability of photon-cell interaction, effectively lowering the scattering and absorbance coefficient (54-56). As a result, more photons were able to penetrate and reach greater depths in the gel-immobilised algal culture. (**Figure 3B**). Such an increase in light penetration depth alleviated self-shading of the algal biomass that is inevitable in dense biofilms, whose thickness is limited to 300 um or less (49, 51, 52), corresponding to an areal biomass density of 30 mm^3^/cm^2^ approximately.

Additionally, gel-immobilised systems were able to achieve significantly higher ***P***_***net***_ compared to biofilm at higher biomass densities, particularly when higher incident irradiance was required to counteract self-shading within the growing biomass (**Figure 3D and 3E**). While photoinhibition could be minimised with a lower incident irradiance, the predominance of a light-limiting regime could lower the overall photosynthetic efficiency, especially in dense biomass with significant self-shading (**Figure 3D**). Hence, as the biomass density increased, moderate to high levels of illumination could reach the shaded cells in the lower region of algal culture. This argument is supported by the decrease in overall ***P***_***net***_ with biomass density and the enhancement of ***P***_***net***_ with incident scalar irradiance beyond areal biomass density of 20 mm^3^/cm^2^ (**Figure 3D**). In comparison to a homogeneous biofilm, a gel-immobilised culture achieved higher ***P***_***net***,_ with an areal biomass density exceeding 30 mm^3^/cm^2^. A heterogeneous biomass distribution from cell aggregation reduced the proportion of photo-inhibited cells among the top layers as not all cells located in the upper layers were exposed to the same intensity as incident illumination. Furthermore, self-shading within individual aggregates protected some of the cells against excessive irradiance. In contrast, the algal biomass in biofilms was uniformly exposed and hence equally photo-inhibited in the top layer. With further self-shading, part of the shielded culture became light-limited (**Figure 3E**). Meanwhile, the shaded biomass beneath the top layer received an optimal light level. In the case of gel-immobilisation, the percentage of photo-inhibited biomass was significantly lower compared to the biofilm, owing to its heterogeneous distribution of cells. Such optimisation required finding an optimal trade-off between alleviating the light shading in the lower region with intense illumination irradiance, while incurring a smaller degree of photoinhibition in the upper region of biomass.

Finally, a gel-immobilised system was able to deliver light more efficiently than a biofilm even when a low incident irradiance is desired, given an optimal scattering matrix. We found that modifying the scattering properties of the gel matrix could enhance the overall photosynthetic performance, both in simulation (**Figure 4C**) and experimentally (**Figure 4E**). These hydrogels showed that algal growth was promoted through increasing the amount of light available to them for photosynthesis. This is especially prominent given the light level that these cultures were grown under was low (∼40 μmol m^-2^ s^-1^). If a higher light level had been used, light may not have been a limiting factor for growth, especially in a growth chamber with direct light falling on the samples. In industrial settings, direct and intense light sources cannot always be guaranteed, and factors like shading or variable sunlight can dilute the light reaching the samples. With the proof of concept shown here, it is now possible to develop a light-sensitive material with dynamic scattering properties upon light exposure of varying intensity, optimising the light distribution with a better trade-off between the proportion of photo-limited and photo-inhibited cells within the population.

However, it is important to take into account that light management is not the sole factor to consider in a photobioreactor. The availability of gases and nutrients in a hydrogel system depends on the diffusion of molecules within and between cell aggregates. Diffusion-limited growth became evident in the lower cell density within a thicker bulk hydrogel (**Figure 4E**). Potential strategies have been studied to address diffusion limitation in hydrogels, such as 3D bioprinting to increase the surface area-to-volume ratio (23, 57) or co-cultivation of algae with symbiotic bacteria to enhance gas and nutrient exchange (39). It should also be noted that this study has assumed a simplified schematic to capture the main optical properties of a homogeneous biofilm. In reality, biofilms growing on a substrate can exhibit more complex morphology both in terms of uneven surface morphology and bulk porosity (49, 58).

In conclusion, our study demonstrates the advantages of cell aggregation in gel-encapsulated colonies of *C. reinhardtii* compared to biofilm growth. This aggregation led to improved light transmission and utilisation, particularly under optimal incident irradiance. As biomass density increased and self-shading became more prominent, the aggregated system achieved a better balance between photo-limited and photo-inhibited regimes when exposed to higher incident irradiance. Furthermore, the addition of scattering particles enhanced light harvesting efficiency, resulting in increased growth rates of *C. vulgaris* under low incident irradiance. By highlighting the collective improvement in light allocation throughout the gel culture, our findings offer new insights for optimising the design and light use efficiency of photobioreactors and microalgae-based photosynthetic living materials.

## Materials and Methods

### Cell culture of *Chlamydomonas reinhardtii* and *Chlorella vulgaris*

The green alga *Chlamydomonas reinhardtii* (wild type strain 137c) was grown mixotrophically in carbon-supplemented Tris-acetate-phosphate (TAP) medium (59) (Tris base: 48.4 mg L^-1^; Beijernick salts (NH_4_Cl: 375 mg L^-1^, MgSO_4_·7H_2_O: 100 mg L^-1^, and CaCl_2_·2H_2_O: 50 mg L^-1^); phosphate solution (K_2_HPO_4_, KH_2_PO_4_); Kropat’s trace elements; CoCl_2_: 1 mg L^-1^; glacial acetic acid: 0.1 *vol*%). Liquid cultures of *C. reinhardtii* were grown in an orbital incubator (Infors HT Multitron Pro.) at 25°C with shaking at 100 rpm under a diurnal cycle of 12h light (100 μmol·photons m^−2^·s^−1^) and 12h dark. *Chlorella vulgaris*, was cultured in Jaworski’s medium (JM) (60), under these conditions: 16h at 25°C under ∼40 μmol m^-2^ s^-1^ and 8h dark, at 20°C in a Panasonic MLR-352-PE growth chamber, unshaken.

### Gel immobilisation of microalgae

For cell immobilisation, 1% w/v agarose (Sigma-Aldrich A9045) with a low gelling temperature (26-30°C) was adopted as the hydrogel matrix in this work. Exponential growth phase cells were taken from suspension cultures, grown under the same conditions, for inoculation into the hydrogels. All work was performed in a flow bench (Air Science Purair Flow-24) to maintain sterile cultures. Gel cultures of *C. reinhardtii* were prepared with 5 % v/v of microalgal cell suspension mixed with agarose solution at varied inoculum densities ranging between 0.1 and 1 million cells per mL prior to gel solidification at 30°C. A 400 μL aliquot of the mixture was allowed to set into a disc that fully covered the observation area (20 mm in diameter) in a 35mm glass-bottomed μ-dish (Ibidi GmbH, Gräfelfing, Germany). *Chlorella vulgaris* were embedded in an agarose matrix, with an inoculum density of 7 million cells per mL. Different thicknesses (1 mm, 2 mm) of agarose (1% w/v) were fabricated in 35 mm glass-bottomed μ-dishes, allowing for *in situ* optical characterisation while maintaining a sterile environment within the petri dish. Within the petri dish, the hydrogels were submerged under 2 mL of liquid JM, that was replenished approximately weekly.

### Cellulose microparticles

To enhance scattering, CMPs (1% w/v) suspended in JM were used, added in varying amounts on a w/v % basis to create agarose with different densities of CMP (0% w/v, 0.2% w/v, 0.5% w/v, 0.7% w/v). Two different solutions were prepared to allow for different final concentrations of CMP in the agarose. A: standard JM and B: the JM with 1% CMP w/v mixture. These solutions were combined in appropriate volumes, e.g., 80% A and 20% B for a 0.2% w/v CMP hydrogel, and this was then used as the precursor mix in which the agarose was dissolved. The precursor plus agarose mixture was then autoclaved to sterilise and initiate crosslinking. The CMPs were isolated from microcrystalline cellulose (SERVA Electrophoresis), with dimensions of approximately 520 nm by 2700 nm. All concentrations of CMP were used for the 1 mm thick hydrogels, while only 0%w/v, 0.5%w/v and 0.7%w/v were used with 2 mm thick gels. Gas diffusion was increasingly problematic as the thickness of the hydrogel increased, causing air bubbles to form within the matrix as the biomass grew.

### Optical density measurements

Optical density (OD) measurements were recorded using an integrating sphere (Labsphere) connected to a spectrometer (Avantes AvaSpec-ULS-RS-TEC) and light source (ThorLabs SLS201L/M), coupled via two 1 mm core fibres (FC-1000-2-SR, Avantes) to the sphere from the light source, and from the sphere to the spectrometer. Normalised transmission measurements were taken as a means to record OD, as there was negligible reflection. All measurements were taken in a dark room. The background signal was acquired with an agarose hydrogel, without algae and the corresponding amount of CMP, also covered in 2 mL of JM, without illumination. The reference signal was taken with the same configuration, however, with the illumination light on. Before measurements, the light source was left on for an hour to allow for it to stabilise. A measurement of the haze of the hydrogels was taken to quantify the scattering. To measure this, transmission spectra were taken of the hydrogels with CMP added but no *C. vulgaris*. A second measurement in the same configuration was taken but with a port removed from the integrating sphere to allow for any ballistic transmission to pass out of the sphere and not be recorded as part of the spectrum. The difference between these two spectra indicates the amount of light being scattered.

To quantify the biomass growth of immobilised cultures, a standard curve between the OD and cell numbers was established. To take OD measurements, the hydrogels were referenced to agarose hydrogels without algae, with the corresponding proportion of scattering particles, also covered in 2 mL of JM. The normalised transmission was measured and given there was negligible reflection, the normalised absorption was calculated by subtracting the transmission from 1 (e.g., 100% transmission). To extract the biomass, the hydrogels were heated to 65°C for 5 minutes (Grant Bio PHMT-PSC24N Thermo-Shaker), and a further 1 mL of JM was added to prevent agarose crosslinking upon cooling. This suspension was then vortexed (Cleaver Scientific) for 10 minutes to break up cell aggregates and cell numbers were counted using a Neubauer Improved Haemocytometer Counting Chamber.

### Cryogenic scanning electron microscopy (CryoSEM)

CryoSEM images of hydrogels were taken of the 1 mm hydrogels with either no CMP added, or 70% CMP. The CMP scattering centres appeared sheet like among the agarose polymer network. A FEI Verios 460 scanning electron microscope with a Quorum cryo-transfer system PP3010T at 2.0 kV was used. Samples were prepared by quenching in liquid ethane, sublimated at -90°C (2 minutes) and finally sputter coated with platinum (10 nm; Quorum Technologies Q150T ES).

Confocal laser scanning microscopy

The size and organisation of cells in the algal aggregates were imaged with a confocal laser scanning microscope (Leica TCS SP5; inverted DMI 6000 CS microscope base). Using a HeNe laser for excitation at 633 nm, the detection range was set to be 660–710 nm, thus capturing the chlorophyll autofluorescence from *C. reinhardtii* centered at 680 nm. Low magnification imaging with a 10X objective (HC PL Fluotar, NA 0.3, Leica, Germany) was used to characterise algal colony size throughout the agarose hydrogel matrix volume by performing a z-stack across varied depth levels from the glass bottom interface to the top of the gel surface. For each sample, z-stack scanning was performed with tile stitching across a lateral area of 4 mm by 4 mm. Individual algal colonies within the immobilisation matrix were also imaged using a 40X oil immersion objective (HCX PL APO CS, NA 1.25, Leica, Germany).

### Optical characterisation of algal aggregates

We imaged individual algal aggregates confined within an agarose gel matrix using a commercially available optical coherence tomography system (Ganymede II, Thorlabs GmbH, Dachau, Germany) equipped with a broadband low coherent superluminescent diode (SLD) emitting a light beam centered at 930nm, and an objective lens with an effective focal length of 18 mm and a working distance of 7.5 mm (LSM02-BB; Thorlabs GmbH, Dachau, Germany). The experimental details followed that of a previous study (61). We evaluated μ_*s*_and *g* within algal aggregates by measuring and fitting the reflectance signal *R* to an exponential decay equation (62-64).

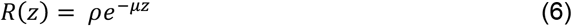

where ρ refers to reflectivity (unitless) at the aggregate interface while *μ* represents the attenuation of light (reciprocal length).

The experimental set-up for microscale light measurements is illustrated in the schematics (**Figure 1G**). Isolated colonies of *C. reinhardtii* with very low inoculum density were grown in 1%w/v agarose gel enclosed within a 35mm glass-bottomed μ-dish. The observation dish was illuminated from the bottom by a fibre-optic halogen lamp (Schott KL-2500, Mainz, Germany) with controlled intensity. Approaching from the top, a fibre-optic microprobe with a spherical tip diameter of 40 μm (60) was used to measure depth profiles of spectral scalar irradiance in vertical steps of 50 μm at specific positions across immobilised algal aggregates (65-67). The position of the probe tip was controlled precisely using a motorised micromanipulator system controlled by custom-built software (68).

Total transmittance spectra of gel-immobilised aggregates were measured using an integrating sphere (Labsphere inc., USA). The illumination port of the integrating sphere was coupled to a stabilised Tungsten-Halogen light source (SLS201L/M, Thorlabs, USA) *via* a 1-mm optical fibre, giving rise to a circular illuminating beam diameter of approximately 5 mm. Similarly, a 1-mm optical fibre was used to couple the detector port of the integrating sphere to a fibre-optic spectrometer (AvaSpec-HS2048, Avantes, USA). The transmitted light intensity was normalised with respect to an agarose gel pad without algae. The spectral measurements were performed by collecting light over 1 s and averaging over three measurements cycles for each spectrum with background subtraction. All measurements were conducted as triplicates.

### Oxygen microsensor measurements

We measured the O_2_ concentration profiles across gel-immobilised *C. reinhardtii* aggregates with a Clark-type O_2_ microsensor (tip size of 25 μm, 90% response time of <0.5 s, and a stirring sensitivity of ∼1%; Unisense A/S, Aarhus, Denmark), as described previously (69). Linear calibration of the sensor readings was performed in an air-saturated and anoxic nutrient medium used for algal cultivation. The percentage of air saturation was converted to absolute oxygen concentration (μmol O_2_ L^-1^) using tabulated values of O_2_ solubility in water as a function of temperature and salinity (Ramsing and Gundersen, Unisense, Denmark; www.unisense.com) depending on the experimental temperature and salinity. All O_2_ microsensor measurements were performed in the same spatial configuration as the scalar irradiance microsensor measurements, with precise positional control using the manipulator at a smaller step interval of 25 μm. Every time the illumination state was changed, we waited for five to ten minutes to attain the steady-state of oxygen profile. The flux of oxygen production and consumption *J* (mol·cm^-2^ s^−1^) were determined from the slopes of oxygen profile using Fick’s first law of diffusion:

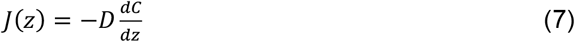

where *D* is the effective diffusion coefficient (cm^2^ s^−1^) while *C* indicates the oxygen concentration at specific positions (mol·cm^-3^). The sum of the flux out of the isolated aggregate was considered the net oxygen production and hence the net photosynthetic rate ***P***_***net***_.

### Three-dimensional voxel-based Monte Carlo simulation of photon transport

We adapted the three-dimensional Monte Carlo simulation program *mcxyz* (41) for our modelling of radiative transfer. The simulation began by “launching” photons sequentially with an equal starting weight, which then propagated with a step size *s* that was determined stochastically. By tracking the photon weight accumulated at individual voxels depending on the optical properties, the light distribution could be simulated accordingly. The numerical approach of Monte Carlo required an accurate input of wavelength-specific optical parameters including refractive index *n*, scattering coefficient μ_*s*_, absorption coefficient μ_*a*,_ and the anisotropy factor *g* of both the gel matrix and encapsulated microalgae. The optical parameters determined experimentally were fed into the *mcxyz* for the simulation of light propagation within different configurations. It allowed the creation of a 3D Cartesian grid of voxels with heterogeneous distribution of media. The input file was first created *via* MATLAB to specify the simulation volume in terms of the number of bins and bin size, and the spatial arrangement of different medium types. The output data would then be generated as a 3D array holding the spatial distribution of normalised scalar irradiance. Absorbance per unit volume per incident energy is the product of normalised scalar irradiance and μ_*a*_. All simulated outcomes were compiled from an average of at least five separate runs, to account for the uncertainty in the random distribution of algal aggregates within the simulated volume.

## Acknowledgments

This work was supported by the ERC BiTe ERC*-*2020*-*CoS-101001637 to S.V. A.S and S-T.C., the Harding Distinguished Postgraduate Scholarship to S.T.C., the Biotechnology and Biological Sciences Research Council (BB/M011194/1) to A.S., EU Horizon 2020 program (H2020-MSCA-ITN-2019) grant N 860125 ‘BEEP to M.M., the Independent Research Fund Denmark (DFF-8022-00301B & DFF-8021-00308B), the Gordon and Betty Moore Foundation (grant no. GBMF9206; https://doi.org/10.37807/GBMF9206) to M.K., and the Gordon and Betty Moore Foundation (GBMF9325) to D.W. and the National Science Foundation (NSF-IntBIO, Award #2316391) to D.W. We acknowledge support from the Swiss National Foundation (Sinergia #198750). We would like to thank the Culture Collection of Algae and Protozoa (CCAP) for providing the wild-type microalgae strain 137c of *Chlamydomonas reinhardtii*. We thank Dr. Yu Kui and Dr. Gianni Jacucci for the insightful discussion and constructive feedback.

