## Supplementary information for "Light management by algal aggregates in living photosynthetic hydrogels"

### Supplementary Information for
Light management by algal aggregates in living photosynthetic hydrogels.

Sing Teng Chua[1*], Alyssa Smith[1*], Swathi Murthy[2], Maria Murace[1], Han Yang[3], Michael Kühl[2], Pietro Cicuta[4], Alison G. Smith[5], Daniel Wangpraseurt[6,7], Silvia Vignolini[1,8]

[1]Yusuf Hamied Department of Chemistry, University of Cambridge, Cambridge CB2 1TN, United Kingdom
[2]Marine Biology Section, Department of Biology, University of Copenhagen, Strandpromenaden 5, DK-3000 Helsingør, Denmark
[3]School of Chemical Engineering, University of Chinese Academy of Sciences, 100040 Beijing, Beijing, China
[4]Cavendish Laboratory, University of Cambridge, CB3 0HE Cambridge, United Kingdom
[5]Department of Plant Sciences, University of Cambridge, Cambridge CB2 3EA, United Kingdom
[6]Marine Biology Research Division, Scripps Institution of Oceanography, University of California San Diego, La Jolla, CA 92093-0205, USA
[7]Department of Nanoengineering, University of California San Diego, La Jolla, CA 92093-0205, USA
[8]Max Planck Institute of Colloids and Interfaces, 14476 Potsdam, Germany

Correspondence: Silvia Vignolini
Co-correspondence: Daniel Wangpraseurt, Sing Teng Chua, Alyssa Smith


**This PDF file includes:**

Figures S1 to S9

**Extraction of optical parameters using OCT**

Optical properties namely, the scattering coefficient $\mu_s$ [$cm^{-1}$] and the anisotropy of scattering *g* for the gel immobilized algal aggregates was obtained using theoretical models of light propagation based on the inverse Monte Carlo method (1). A more detailed description of extraction of optical properties from OCT scans can be found elsewhere (2, 3).

Briefly, OCT B-scans were acquired with a resolution of 581×1024 pixels, over a fixed depth of 2.8 mm, and variable distance in the X-plane. The setup was optimized to yield the highest signal at a fixed distance of 0.4 mm from the top of the scan. The OCT reflectivity (R) was calibrated (**SI Figure 2A**) before measurements using homemade reflectance standards with an immersion oil-glass, a water-glass, and an air-glass interface. R values from the standards were determined using Fresnel's equation:

Eq. 1: $$R = \left(\frac{n_1 - n_2}{n_1 + n_2}\right)^2$$

using the refractive index (n) for air (1), water (1.33), immersion oil (1.46), and quartz glass (1.52). The OCT signal (in decibel, dB), from the samples, was then converted to the depth-dependent R via a linear fit of $\log_{10}(R)$ versus OCT intensity values (see reference (3) for details).

The focus function of the objective lens was calibrated by measuring the OCT signal fall off, in steps of 0.1 mm, from either side of the focal plane (z = 0.4 mm) to z = 0 and 0.8, respectively. The signal loss from the focal plane follows an exponential decay function. The determined R values from the sample scans were corrected by dividing with the exponential fit. The corrected R values were then plotted against sample depth (z, distance from focal volume) and fitted to the exponential decay function (**SI Figure 2B**):

Eq. 2: $$R(z) = \rho \times e^{-\mu . z}$$

where ρ (dimensionless) is the light intensity and μ is the signal attenuation ($cm^{-1}$) from the focal volume. The fit was considered satisfactory if $R^2 > 0.5$.

Using the grid method (4), values of ρ and μ were mapped to *g* and $\mu_s$ based on the theory described in other previous studies (3, 5). It was assumed that the sample absorption at 930 nm was negligible and the absorption was dominated by water ($\mu_a$ – 0.43 $cm^{-1}$). The effective numerical aperture (NA) was 0.11.

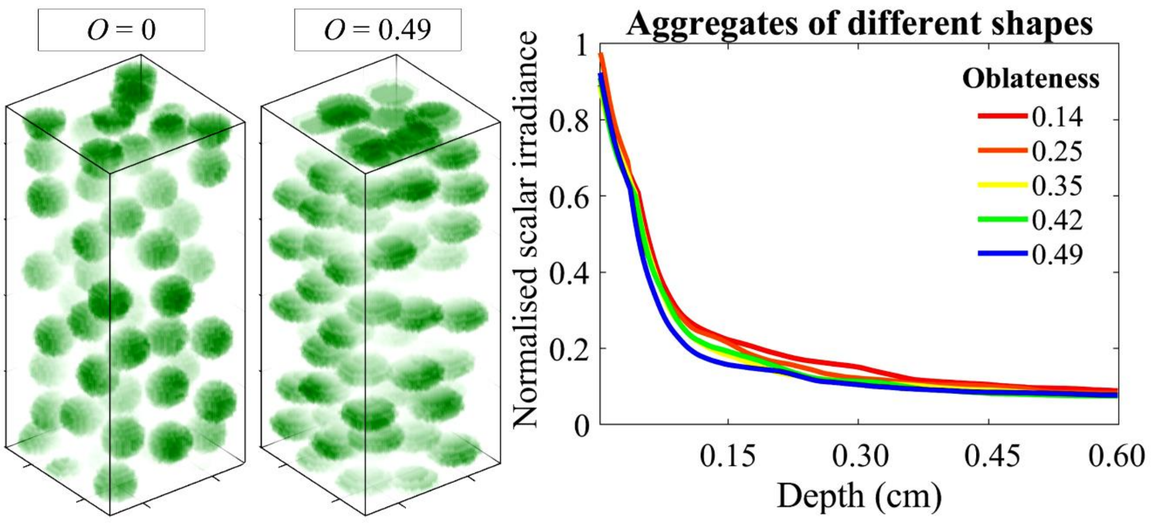


**Fig. S1.** Simulation of fluence attenuation among microalgal aggregates of different oblateness (*O*). $O = \frac{a-c}{a}$, where a, c = major, minor axis.

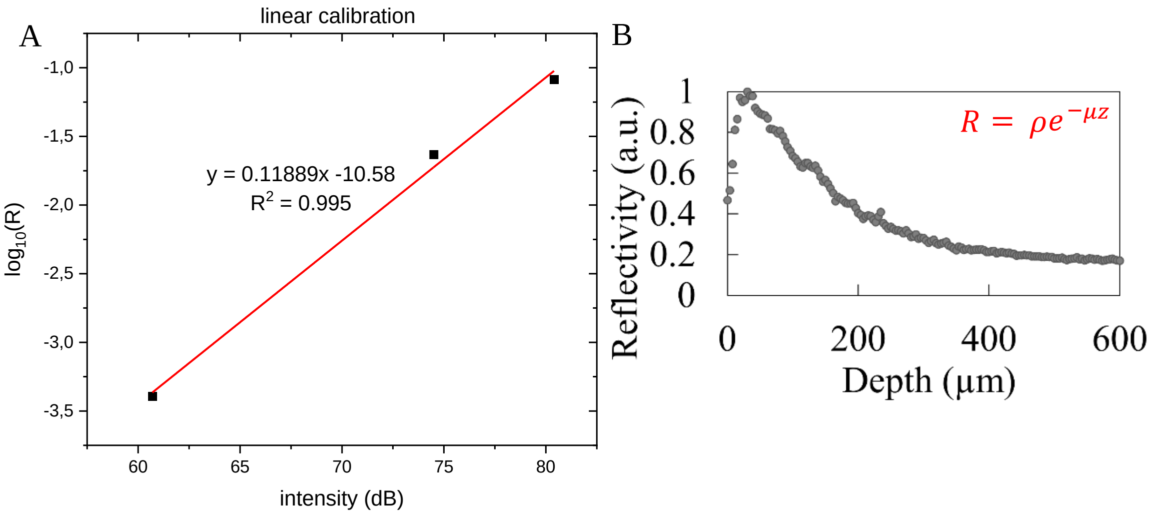


**Fig. S2.** (A) Calibrated reflectivity versus intensity in dB; (B) Reflectivity profile from an optical coherence tomography (OCT) scan of an algal aggregate illustrating the exponential decay of backscattered signal with depth into the aggregate, which could be fitted with the empirical model (5).

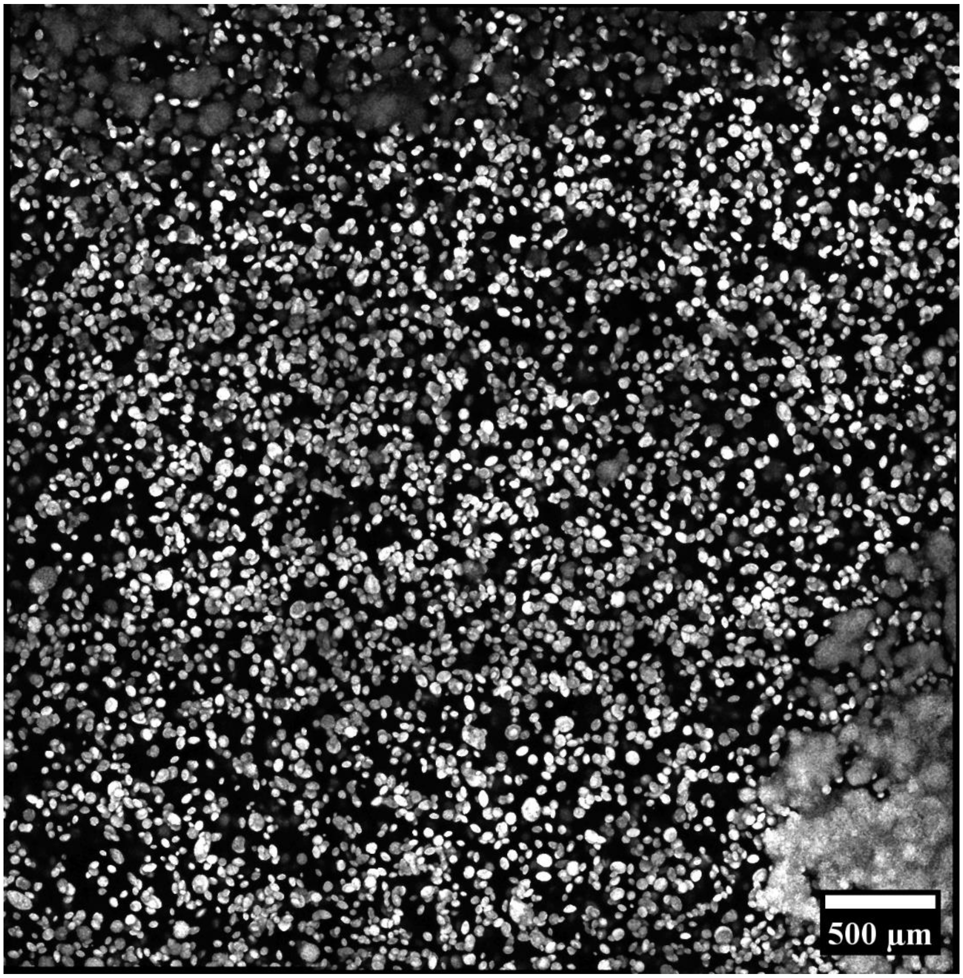


**Fig. S3.** Confocal imaging (maximum Z projection from a 100-um stack) of *Chlorella vulgaris* aggregates after 7 days of growth within an agarose hydrogel.

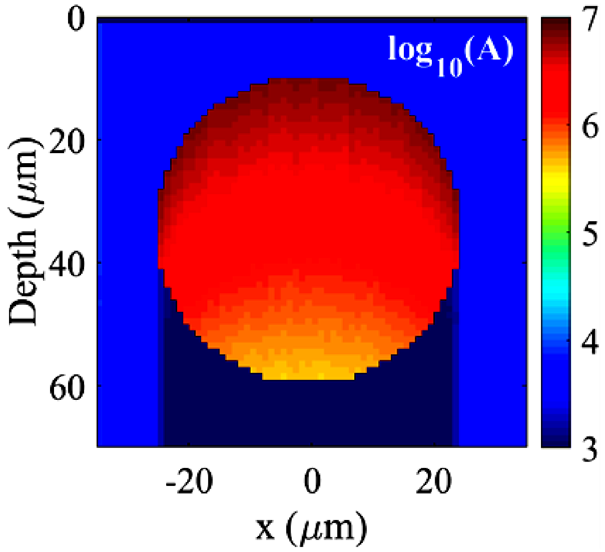


**Fig. S4.** Monte Carlo simulation of light attenuation within a single aggregate (cross-sectional view). The colour bar represents light absorbance in logarithmic scale and in arbitrary units.

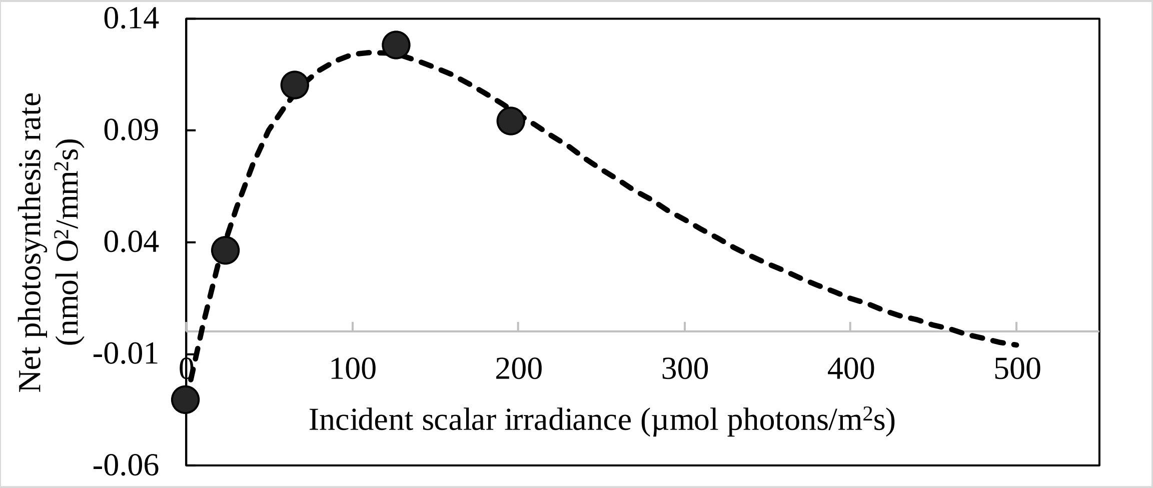


**Fig. S5.** Light response curve obtained experimentally with microsensor measurements of scalar irradiance and oxygen concentration within an isolate algal aggregate, fitted to the empirical model of Platt et al.. (6).

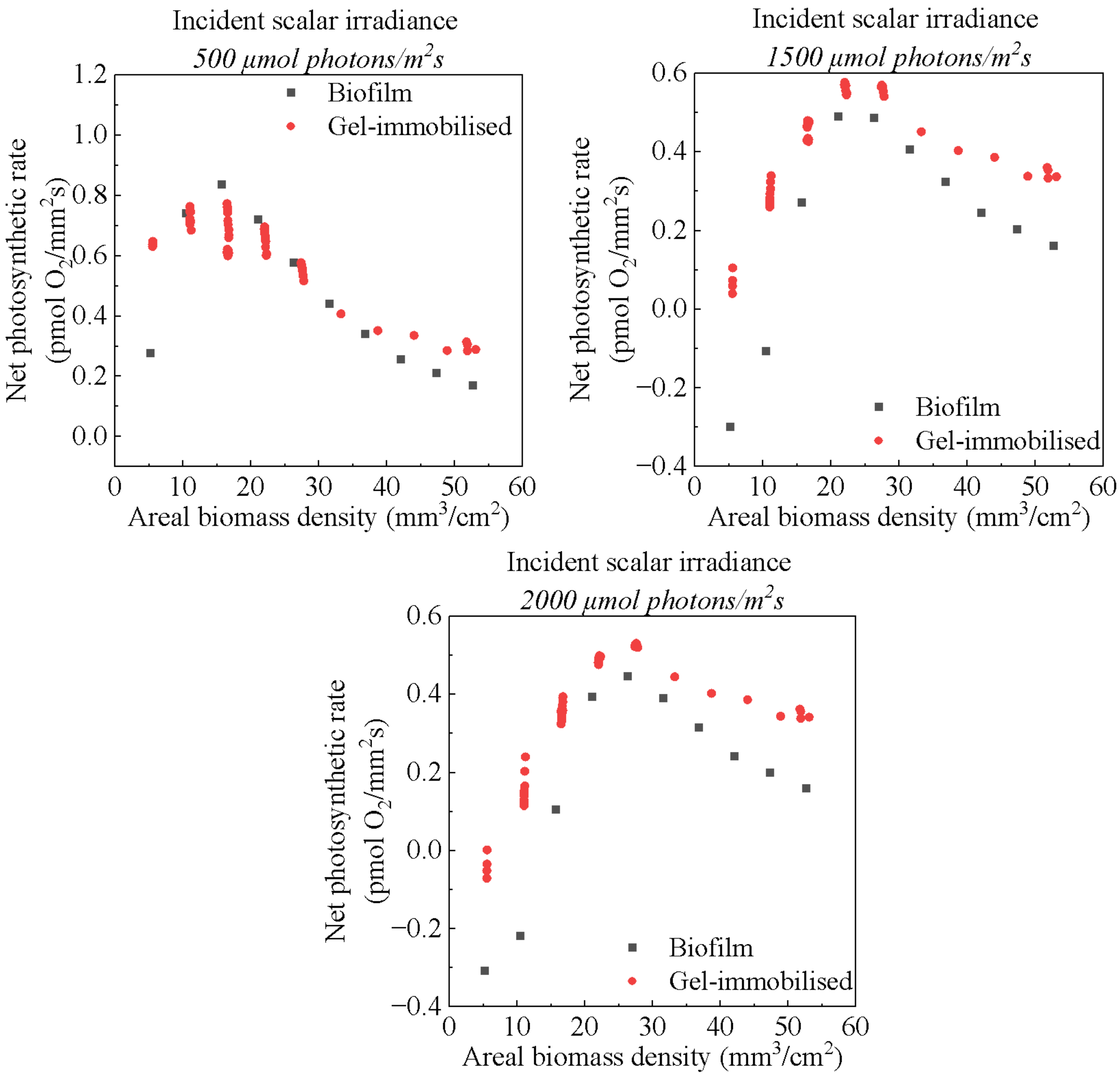


**Fig. S6.** Calculation of the net photosynthetic rate by coupling the experimental light response curve to the simulated fluence variation among algal aggregates and through biofilm of different areal biomass densities under an incident scalar irradiance of 500, 1500 and 2000 µmol photons/$m^2$s respectively.

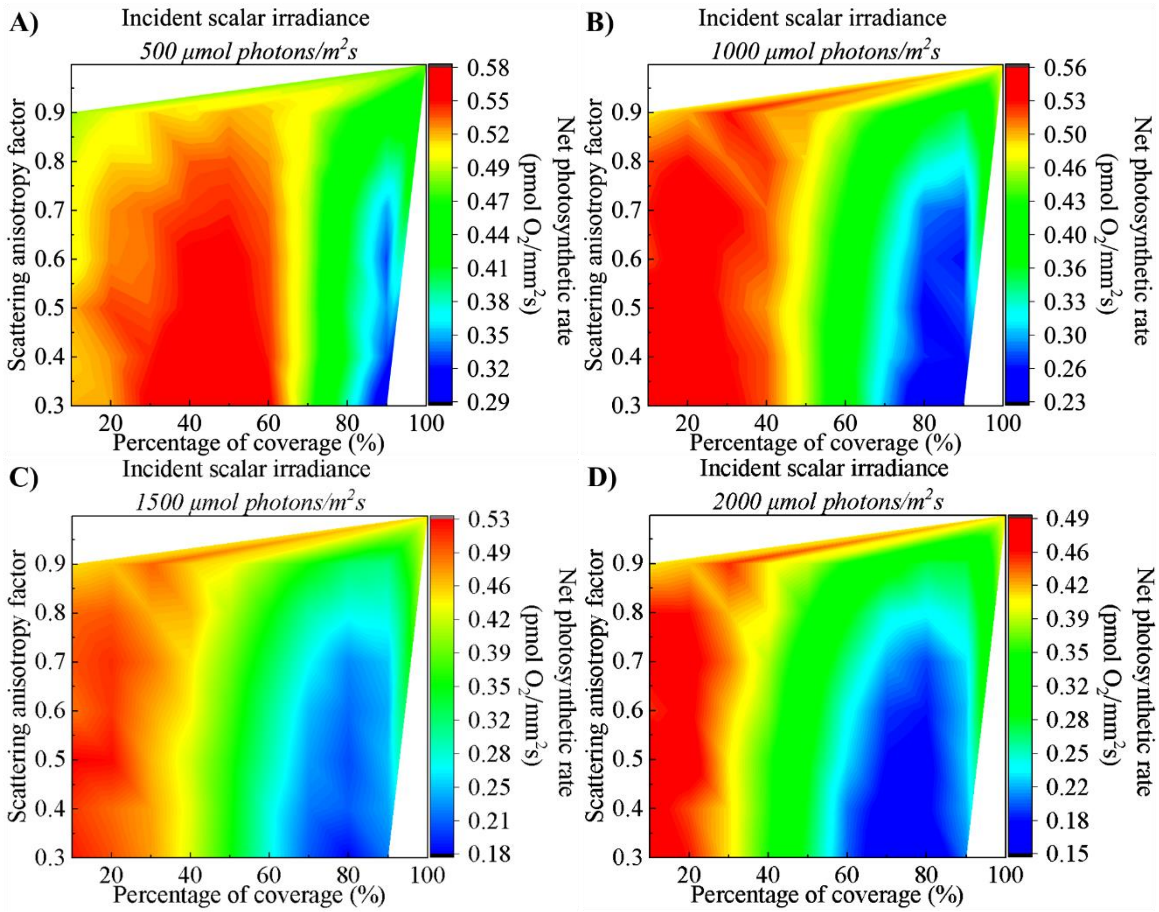


**Fig. S7.** Calculation of the net photosynthetic rate by coupling the experimental light response curve to the simulated variation of normalised scalar irradiance among algal aggregates within different scattering matrix configurations under an incident scalar irradiance of (A) 500 µmol photons $m^{-2}$ $s^{-1}$**;** (B) 1000 µmol photons $m^{-2}$ $s^{-1}$**;** (C) 1500 µmol photons $m^{-2}$ $s^{-1}$**;** (D) 2000 µmol photons $m^{-2}$ $s^{-1}$.

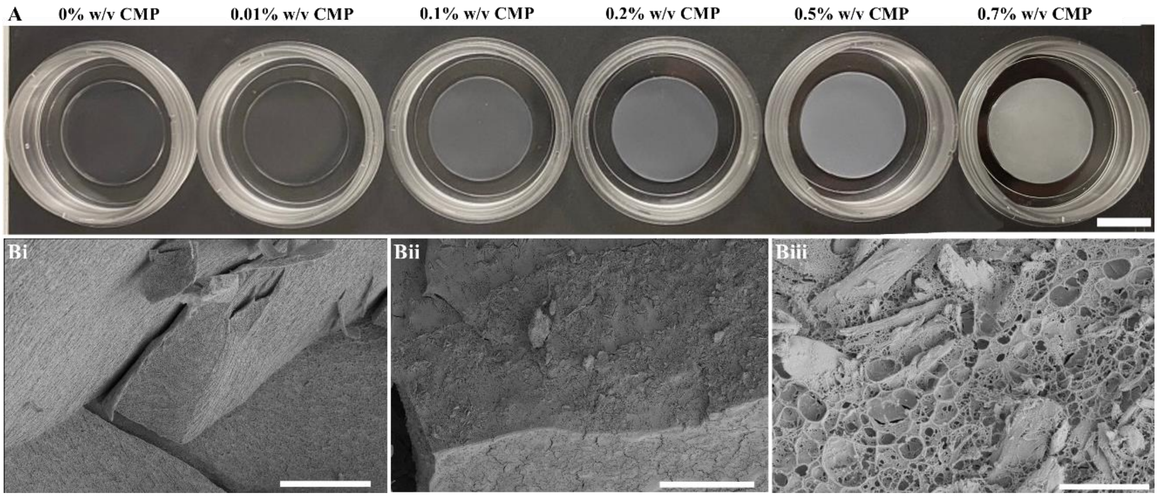


**Fig. S8.** (A) Visual effect of cellulose microparticles (CMP) doping on agarose gel pads, with the weight percentage of CMP doping shown above the image. Scale bar = 1 mm. (B) Cryogenic scanning electron microscope images of hydrogels: (i) 0%w/v CMP (Scale bar = 20 µm); (ii) 0.7%w/v CMP (Scale bar = 20 µm) and (iii) 0.7%w/v CMP (Scale bar = 5 µm). The CMP scattering centres appear sheet-like among the agarose networks.

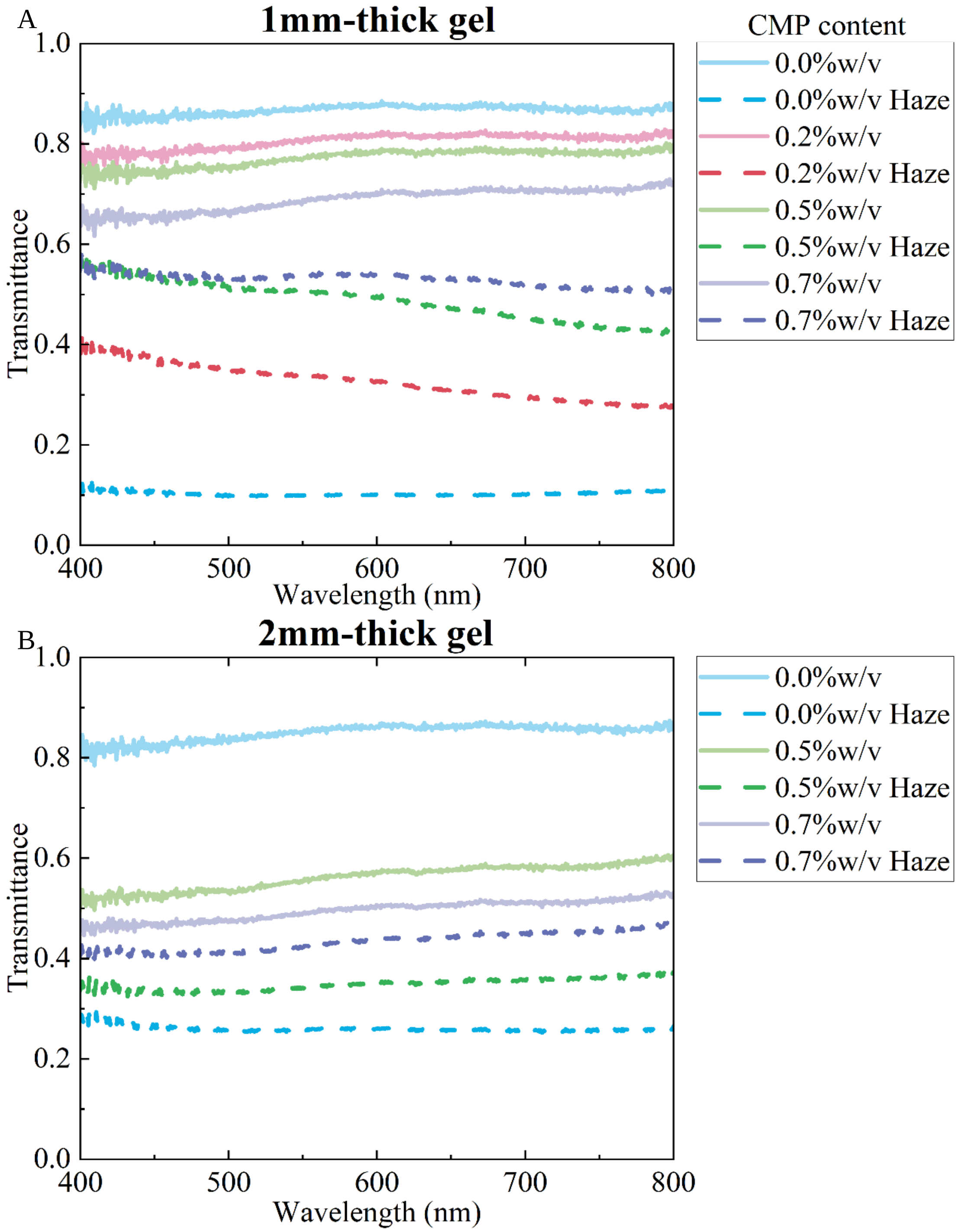


**Fig. S9.** The effect of cellulose microparticles (CMP) doping on the transmission and scattering within 1 mm (A) and 2 mm (B) thick agarose hydrogels, across different levels of embedded CMP. The transmission can be seen with the solid line and the haze spectra is shown with the dashed line.
